# The reported healthy ageing gene expression score: lack of association in two cohorts

**DOI:** 10.1101/034058

**Authors:** Luke C. Pilling, Lorna W Harries, Dena G Hernandez, Andrew B Singleton, George A. Kuchel, Luigi Ferrucci, David Melzer

## Abstract

Sood *et al*. report a multi-tissue RNA signature “predictive of human health, using only peripheral blood samples”. We tested this score in blood in two independent, larger cohorts and found no associations with age or related phenotypes, including muscle strength, interleukin-6 or mortality.

## Correspondence

Sood *et al*. (1) report identifying a 150-gene classifier from muscle samples from 15 ‘healthy’ older (59 to 77 years old) and 15 ‘healthy’ younger (19 to 28 years) subjects (1), and presented this as an “RNA signature of human healthy ageing that can act as a diagnostic of future health, using only a peripheral blood sample”. Sood *et al*. tested the classifier in muscle, brain and skin expression data (up to age 86), with reported success in classifying young versus old. Using data from the ULSAM cohort (males aged 70 years), the classifier was collapsed into a “healthy ageing gene score” (HAGS), which was tested against clinical parameters in linear models, in which HAGS explained 11% of the variance in renal function 12 years later (measured by cystatin C, Sood *et al*. table S1). An association with mortality was also claimed (unadjusted p=0.03), but statistical tests were noted for 26 clinical measures, two diseases plus mortality in this sample of only 108 men, with no adjustment for multiple statistical testing.

Evidence for HAGS in blood samples was limited to Sood *et al*. classification performance in patients already suffering from Alzheimer’s disease (AD), mild cognitive impairment (MCI) and controls (with no data on future ‘health’). HAGS performance alone in classifying AD vs controls was modest (ROC areas under the curve 66% and 73% in two datasets: 50%=random, 100%=perfect ‘diagnosis’) and HAGS alone could not distinguish between MCI and AD. No other data were presented on HAGS as a ‘diagnostic of future health’ in blood or other tissues.

Here we investigated HAGS using whole blood-derived expression data from two well-characterized human cohorts including wide age ranges and both sexes; the Italian aging cohort, InCHIANTI (2), and the Baltimore Longitudinal Study of Aging, BLSA (3). We hypothesized that a gene expression ageing score should be positively correlated with chronological age in linear modelling and should predict mortality. It should also be Pilling *et al*. – Healthy Ageing gene expression score in two human cohorts associated with commonly studied age-related phenotypes including muscle strength, cognitive, renal function and inflammation.

Sood *et al*. report applying the 150 Affymetrix array derived gene set in data from Illumina Human HT-12 v3 microarrays used in skin and blood samples, with 128 genes included. In our Illumina HT-12 based data, we found that only 119 (of the 150) gene IDs had a corresponding probe in the Illumina Human HT-12 v3 array annotation (4). Some genes mapped to more than one probe; we selected the probe with the highest mean expression across the participants for each gene. Of these 119 genes, 77 were significantly expressed above background (v3 array, p<0.01 in at least 5% of participants (5)) in InCHIANTI and 66 (v4 array) in BLSA. We therefore calculated associations with both the genes expressed above background and the full n=119 sets separately. We created the HAGS using the methods of Sood *et al*., where the ‘direction’ of each gene’s association with age was determined in Supplementary Table 1 of (1).

Details of the sample collection, quality assurance and analysis of the InCHIANTI data are set out in Harries *et al*. 2011 (5), and similar methods were used in BLSA. Robust linear regression models (STATA v13) against HAGS were adjusted for age, microarray batch and basic white blood cell subtype counts (number of lymphocytes, monocytes, eosinophils and basophils). InCHIANTI was also adjusted for the two study sites and BLSA for race (white/other). Cox proportional hazards survival models were used for all-cause mortality, with the same covariates as above.

Summary statistics for the study participants are provided in Table 1. Maximum follow-up time periods for InCHIANTI and BLSA were approximately 6.7 years and 6 years, respectively. HAGS based on genes expressed above background was not associated with our studied ageing phenotypes (Table 2) in cross-sectional analyses in either cohort. Renal function, estimated by Cockcroft-Gault creatinine clearance, was associated with HAGS in InCHIANTI but this association became non-significant after adjustment for BIA-estimated whole body muscle mass (%), known to influence creatinine clearance estimates in the elderly. There was no association between HAGS and mortality over 6-years, either in a categorical analysis (Table 2) or in a continuous analysis of trend (HR=0.99, p=0.84, and HR=0.99, p=0.55, in InCHIANTI and BLSA respectively). Data were available on Mini Mental State Examination score (MMSE, a widely used cognitive impairment measure) in the InCHIANTI cohort, but we found no associations (p>0.05) with HAGS using linear (mean=25.6, sd=5.6, range=0 to 30) or categorical analyses (MMSE categories: [0]=0/23, [1]=24/28, [2]=29/30).

**Table 1.** Characteristics of the cohorts

|  | InCHIANTI | BLSA |
| --- | --- | --- |
| N | 695 | 763 |
| Age (mean, SD) | 72 (15.3) | 68 (12.9) |
| Age (min, max) | 30 - 104 | 31 - 98 |
| Females (n, %) | 382 (55%) | 368 (48%) |
| Race, white (n, %) | 695 (100%) | 517 (68%) |
| Died during follow-up (n, %) | 156 (23%) | 45 (6%) |
| Years to death since RNA visit (mean, SD) | 3.45 (1.77) | 3.04 (1.48) |
| Years to death since RNA visit (min, max) | 0.23 - 6.66 | 0.25 - 6.02 |
| Healthy Ageing Gene Score (mean, SD) | 351 (65) | 394 (187) |
| Healthy Ageing Gene Score (min, max) | 117 - 549 | 7 - 763 |

**Table 2.**
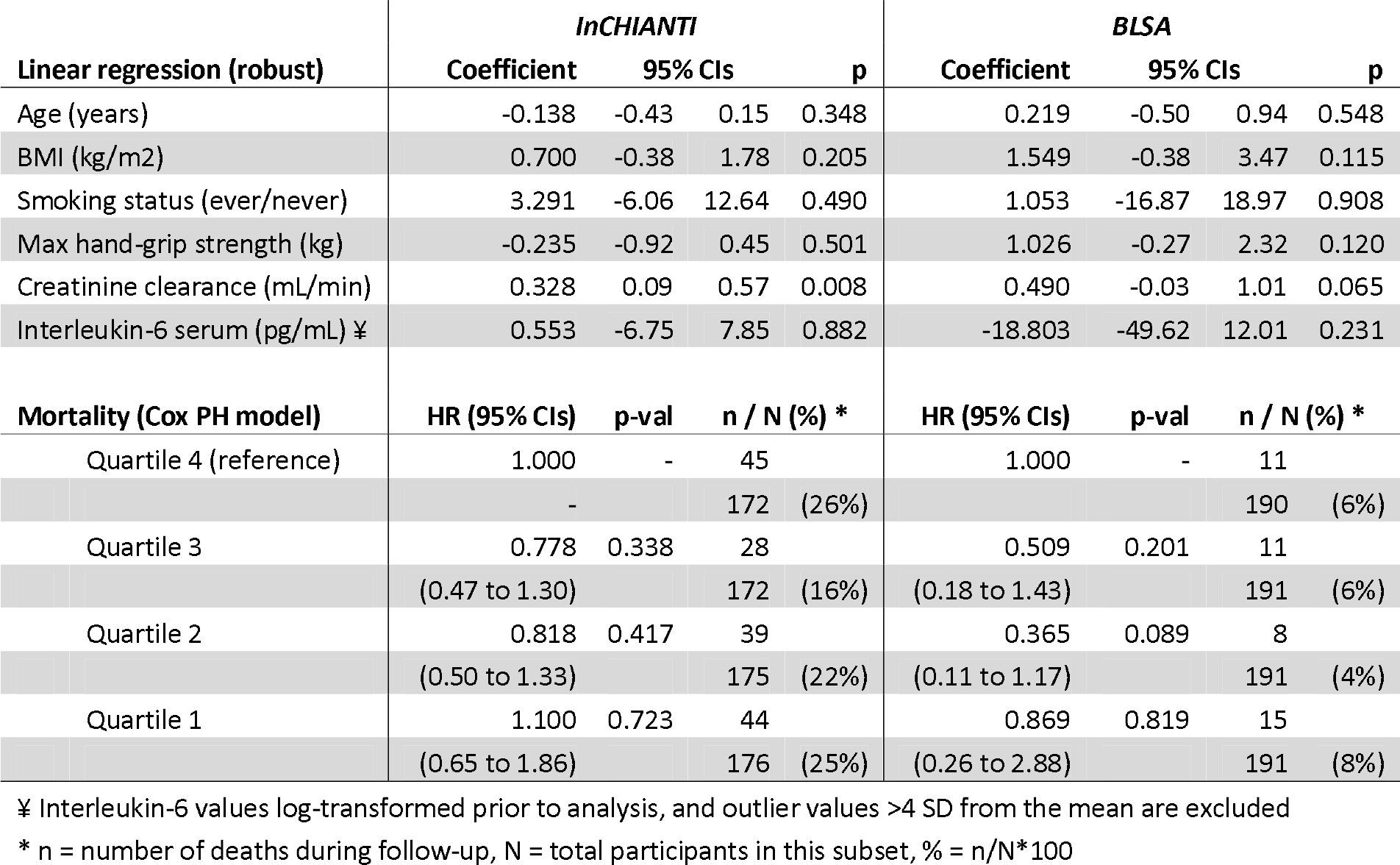
Associations between Healthy Aging Gene Score and selected age-related phenotypes

| Linear regression (robust) | InCHIANTI |  |  |  | BLSA |  |  |  |
| --- | --- | --- | --- | --- | --- | --- | --- | --- |
|  | Coefficient | 95% CIs |  | p | Coefficient | 95% CIs |  | p |
| Age (years) | -0.138 | -0.43 | 0.15 | 0.348 | 0.219 | -0.50 | 0.94 | 0.548 |
| BMI (kg/m<sup>2</sup>) | 0.700 | -0.38 | 1.78 | 0.205 | 1.549 | -0.38 | 3.47 | 0.115 |
| Smoking status (ever/never) | 3.291 | -6.06 | 12.64 | 0.490 | 1.053 | -16.87 | 18.97 | 0.908 |
| Max hand-grip strength (kg) | -0.235 | -0.92 | 0.45 | 0.501 | 1.026 | -0.27 | 2.32 | 0.120 |
| Creatinine clearance (mL/min) | 0.328 | 0.09 | 0.57 | 0.008 | 0.490 | -0.03 | 1.01 | 0.065 |
| Interleukin-6 serum (pg/mL) ¥ | 0.553 | -6.75 | 7.85 | 0.882 | -18.803 | -49.62 | 12.01 | 0.231 |
| Mortality (Cox PH model) | HR (95% CIs) | p-val | n / N (%) * |  | HR (95% CIs) | p-val | n / N (%) * |  |
| Quartile 4 (reference) | 1.000 | - | 45 |  | 1.000 | - | 11 |  |
|  | - |  | 172 (26%) |  |  |  | 190 (6%) |  |
| Quartile 3 | 0.778 | 0.338 | 28 |  | 0.509 | 0.201 | 11 |  |
|  | (0.47 to 1.30) |  | 172 (16%) |  | (0.18 to 1.43) |  | 191 (6%) |  |
| Quartile 2 | 0.818 | 0.417 | 39 |  | 0.365 | 0.089 | 8 |  |
|  | (0.50 to 1.33) |  | 175 (22%) |  | (0.11 to 1.17) |  | 191 (4%) |  |
| Quartile 1 | 1.100 | 0.723 | 44 |  | 0.869 | 0.819 | 15 |  |
|  | (0.65 to 1.86) |  | 176 (25%) |  | (0.26 to 2.88) |  | 191 (8%) |  |

¥ Interleukin-6 values log-transformed prior to analysis, and outlier values >4 SD from the mean are excluded
\* n = number of deaths during follow-up, N = total participants in this subset, % = n/N\*100

Timmons has argued elsewhere (biorxiv http://biorxiv.org/content/early/2015/12/09/034058) that HAGS was developed as a non-linear model for ages 20 to 70 years old. In analyses restricted to 55-70 year-olds (n=82 in InCHIANTI, n=295 BLSA), we re-created the HAGS score and found no associations (p>0.05) with the studied age-related traits, or with mortality (n=6 deaths in InCHIANTI, n=8 in BLSA).

In analyses including all available Illumina HT12 v3 HAGS gene probes (i.e. not excluding probes expressed below background levels), but results were similar. Results were also similar when only including male participants in analyses. Finally, Sood *et al*. do not report adjusting for age as their cohorts were all age-matched; the only difference in our results when not including age (or cell counts, which are correlated with age) as adjustments was that creatinine clearance was no longer associated with HAGS (coef=0.086, p=0.31) in models without adjustment for muscle mass.

In a recent meta-analysis of nearly 15,000 participants from 14 cohorts including InCHIANTI, Peters *et al*. report that expression of 1,497 genes in human whole blood samples had sufficient overall correlations with chronological age to produce significant estimates in linear modelling, with 25% of these also associated with age in brain tissue (6). Of the 150 genes identified by Sood *et al*. only 10 genes were also found to be associated with chronological age in this large-scale meta-analysis in human blood.

We have shown in two relatively large independent cohorts that the recently reported “healthy aging gene score”, measured in blood, was not associated with chronological age or several age-related phenotypes, and was not predictive of mortality over 6 years. These results were consistent across sensitivity analyses using different data subsets (e.g. narrower age ranges) and adjustments.

## Declarations

All authors have read the manuscript and provided input, and declare no competing interests.

